# A stem-loop RNA RIG-I agonist confers prophylactic and therapeutic protection against acute and chronic SARS-CoV-2 infection in mice

**DOI:** 10.1101/2021.06.16.448754

**Authors:** Tianyang Mao, Benjamin Israelow, Carolina Lucas, Chantal B. F. Vogels, Olga Fedorova, Mallery I. Breban, Bridget L. Menasche, Huiping Dong, Melissa Linehan, Yale SARS-CoV-2 Genome Surveillance Initiative, Craig B. Wilen, Marie L. Landry, Nathan D. Grubaugh, Anna M. Pyle, Akiko Iwasaki

**Author notes:** A full list of authors appears at the end of the paper.

## Abstract

As SARS-CoV-2 continues to cause morbidity and mortality around the world, there is an urgent need for the development of effective medical countermeasures. Here, we assessed the antiviral capacity of a minimal RIG-I agonist, stem-loop RNA 14 (SLR14), in viral control, disease prevention, post-infection therapy, and cross-variant protection in mouse models of SARS-CoV-2 infection. A single dose of SLR14 prevented viral replication in the lower respiratory tract and development of severe disease in a type I interferon (IFN-I) dependent manner. SLR14 demonstrated remarkable protective capacity against lethal SARS-CoV-2 infection when used prophylactically and retained considerable efficacy as a therapeutic agent. In immunodeficient mice carrying chronic SARS-CoV-2 infection, SLR14 elicited near-sterilizing innate immunity by inducing IFN-I responses in the absence of the adaptive immune system. In the context of infection with variants of concern (VOC), SLR14 conferred broad protection and uncovered an IFN-I resistance gradient across emerging VOC. These findings demonstrate the therapeutic potential of SLR14 as a host-directed, broad-spectrum antiviral for early post-exposure treatment and for treatment of chronically infected immunosuppressed patients.

---

SARS-CoV-2 is an enveloped, positive-strand RNA virus that causes both upper and lower respiratory infection in humans and other animals^1^. As of May 26^th^ 2021, the ongoing global COVID-19 pandemic caused by SARS-CoV-2 has led to 167.5 million confirmed cases and 3.5 million deaths worldwide, inflicting widespread economic, sociological, and psychological damage. The clinical spectrum of SARS-CoV-2 infection is wide. While most infections are asymptomatic or mild, older patients, significantly those with underlying medical comorbidities and male sex, are more likely to develop severe diseases involving acute respiratory distress syndrome (ARDS), multi-organ failure, and death^2^. Currently, there is a paucity of effective antivirals to treat COVID-19, with remdesivir and monoclonal antibodies demonstrating modest efficacy in a select subset of patients^3,4^. To halt substantial morbidity and mortality from COVID-19 around the globe, in addition to the use of vaccines in preventing the disease, efforts are required to develop efficacious therapeutics against SARS-CoV-2.

Great strides made in the understanding of COVID-19 immunology have provided crucial insights into the central role of IFN-I in host immune responses against SARS-CoV-2 infection^5,6^. The innate immune system utilizes host-encoded nucleic acid sensors, known as the pattern recognition receptors (PRRs), to surveil viral pathogens by detecting their pathogen-associated molecular patterns (PAMPs)^7^. Following SARS-CoV-2 infection, multiple cytosolic PRRs, including RIG-I, MDA-5, and LGP2, mediate viral RNA recognition in infected lung epithelial cells and initiate front-line antiviral defense through the production of IFN-I^8,9^. Upon secretion, IFN-I engages with its universally expressed receptor in autocrine and paracrine fashions, stimulating the expression of a large network of interferon stimulated genes (ISGs) to inhibit viral replication, and cytokines and chemokines to recruit specialized immune cells to sites of infection. In the context of infection with SARS-CoV-2, IFNs appear to play dichotomous roles. While delayed and prolonged type I and III IFNs are associated with severe disease, an early, robust, and regulated production of IFN is protective against COVID-19^10,11^. This is well exemplified by the susceptibility to life-threatening disease of SARS-CoV-2-infected individuals with inborn defects in IFN-I production and signaling or neutralizing autoantibodies against IFN-I^12,13^. COVID-19 patients found with anti-IFN-I autoantibodies demonstrate significantly delayed virological clearance relative to patients without such autoantibodies^14^. In a mouse model of SARS-CoV-2 infection, early IFN-I blockade leads to exacerbation of disease severity^14^. Collectively, these studies highlight the beneficial role of IFN-I in SARS-CoV-2 infection and suggest innate immune sensors as promising therapeutic targets to be harnessed for prevention and treatment of COVID-19.

The innate immune system can be pharmacologically modulated to elicit tailored effector outputs with desired immunological outcomes^15,16^. Given the importance of timely induction of IFN-I in SARS-CoV-2 infection, PRRs can be activated in a targeted manner to induce antiviral protection^17^. Our approach in leveraging a synthetic activator of antiviral immunity to combat SARS-CoV-2 builds on our previous work demonstrating that short, tri- or di-phosphorylated stem-loop RNAs (SLR) act as specific and potent agonists for the cytosolic RNA sensor RIG-I^18,19^. SLRs are designed to mimic physiological double-stranded RNA (dsRNA) ligands for RIG-I by stably folding into a minimal ligand containing 14–base pair RNA duplex (hence the name SLR14) and a tri- or di-phosphorylated 5 ′ terminus. Each SLR14 presents a single duplex terminus and productively binds one RIG-I molecule. The opposite end of the duplex is blocked with a stable RNA tetraloop to ensure that the RIG-I-SLR14 interaction is structurally defined and resistant to nucleases and strand dissociation. Unlike polyinosinic:polycytidylic acid (poly I:C), which is a widely used dsRNA ligand of unknown structure recognized by a handful of PRRs, SLR14 specifically activates RIG-I and triggers IFN-I dominant innate immune response (over IFN-III response) within 2 hours of intravenous injection in mice^19^. With its synthetic simplicity, chemically defined composition, targeted receptor binding, and *in vivo* potency, SLR14 holds great promise as a new class of RNA therapeutics that can be applied as antivirals against SARS-CoV-2.

While most individuals effectively clear SARS-CoV-2 infection, growing evidence suggests that infection in immunocompromised patients, such as those with severe forms of B cell and antibody deficiency, can become chronic^20,21^. In these patients, persistent infection can also foster continuous intra-host viral evolution and lead to further emergence of immune-evasive variants, likely as a result of selective pressure driven by insufficient natural or transferred antibodies. While some patient case reports have used convalescent plasma to treat chronic SARS-CoV-2 infection, currently there are no approved therapeutic options^22^. Although vaccines currently approved against SARS-CoV-2 are effective at preventing severe disease and death in individuals with an intact immune system, their immunogenicity is significantly attenuated in immunocompromised patients, eliciting suboptimal humoral immune responses^23^. Therefore, therapeutic strategies that exert strong antiviral effect independent of adaptive immunity in the setting of immunosuppression are in dire need.

Since the initial outbreak, several SARS-CoV-2 variants have rapidly emerged with enhanced transmissibility and altered immunogenicity. The B.1.1.7 variant is more transmissible than other variants (∼50% increase) and is spreading rapidly around the globe^24,25^. While mutations accumulated by B.1.1.7 seem to have negligible impact on infection- and vaccine-induced antibody immunity^26,27^, other variants have been found to acquire mutations on their spike proteins that can evade antibody targeting. Notably, variant B.1.351 and P.1 have both demonstrated considerable resistance to antibody binding and neutralization^28,29^. The more recent variant B.1.526 has also exhibited some level of antibody evasion^30^. In addition, these variants harbor mutations outside the spike protein that may enable strong antagonism of the host antiviral innate immunity. In this context, the containment of COVID-19 will require prophylactic and therapeutic antiviral strategies that afford cross-variant protection.

## A single dose of SLR14 confers potent antiviral protection against lethal SARS-CoV-2 infection

To examine the antiviral activity of SLR14 *in vivo*, we used a mouse model of SARS-CoV-2 infection that transgenically expresses human angiotensin-converting enzyme 2 (ACE2) under the keratin 18 gene promoter, also known as the K18-hACE2 mice^31^. Intranasal infection with SARS-CoV-2 in K18-hACE2 mice leads to viral replication, pulmonary inflammation, and respiratory dysfunction, recapitulating key aspects of infection and pathogenesis seen in patients with COVID-19^32,33^. We have previously shown that intravenous (i.v.) injection of SLR14 complexed with polyethyleneimine (PEI) results in rapid, short-lived, and systemic IFN-I responses that peak as early as 2 hours post injection and decline to undetectable levels within 24 hours of injection^19^. Based on this, we intranasally infected K18-hACE2 mice with the ancestral strain of SARS-CoV-2 (2019n-CoV/USA_WA1/2020), administered SLR14 i.v. 4 hours post infection, and monitored survival and weight loss daily thereafter (**Fig. 1a**). SLR14 treatment considerably prevented weight loss and dramatically improved survival following the infection (**Fig. 1b-c**). In contrast, vehicle-treated mice uniformly lost weight and developed apparent signs of sickness behaviors such as reduced motility and hyporesponsiveness, rapidly succumbing to infection by 8 days post infection (DPI). These results showed that SLR14 effectively alleviates morbidity and reduces mortality, affording protection against lethal SARS-CoV-2 infection *in vivo*.

**Figure 1.**
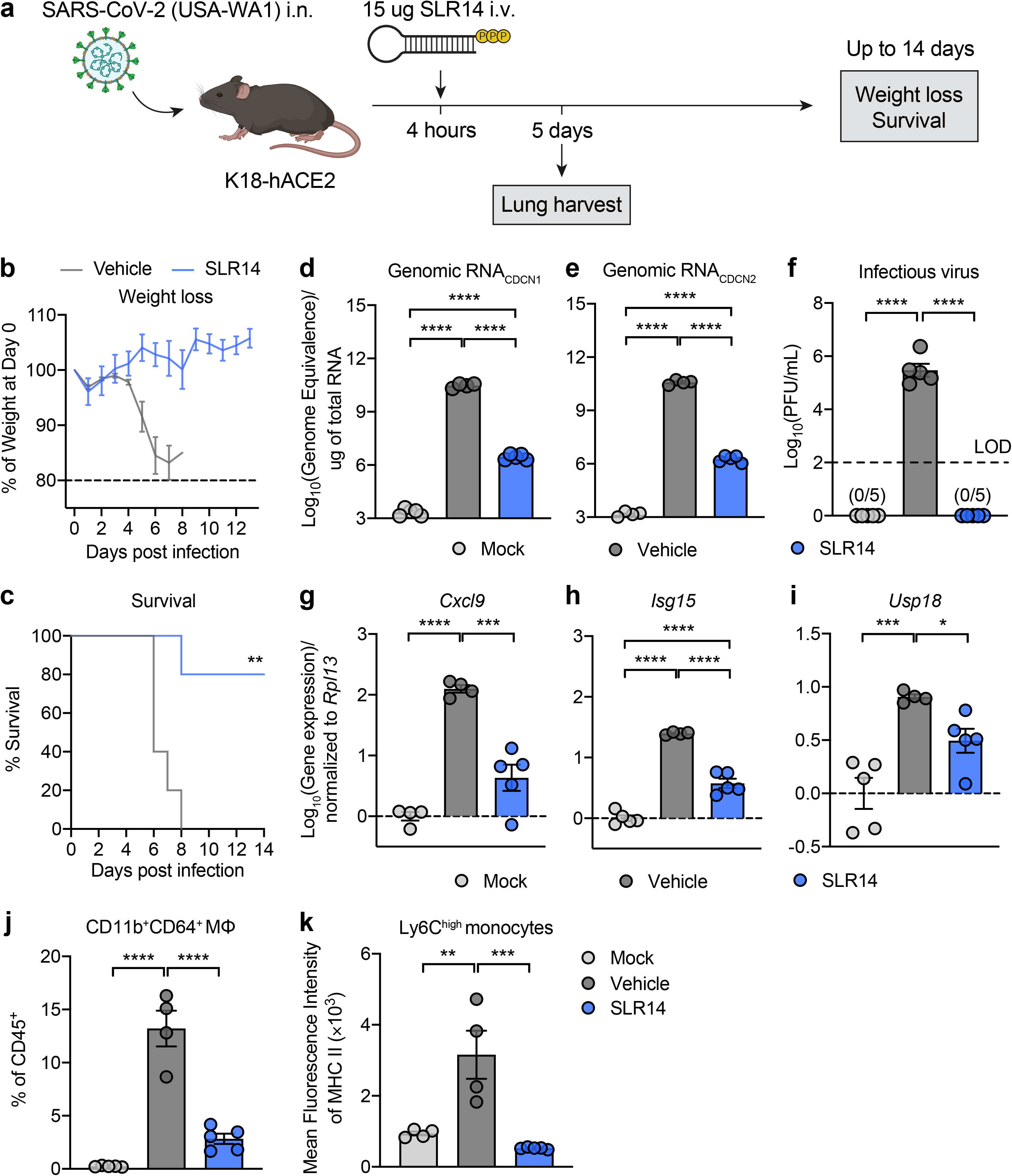
**a**, Experimental scheme: K18-hACE2 mice were intranasally infected with 10^3^ PFU SARS-CoV-2 (2019n-CoV/USA_WA1/2020). 4 hours post infection, 15 μ SLR14 or vehicle were intravenously administered. Weight loss and survival were monitored daily up to 14 DPI. Death was recorded when mice were found dead in the cage, moribund, or at 80% of original body weight. In a separate cohort, lung tissues were collected for virological and immunological analysis 5 DPI. **b,c**, Weight loss (**b**) and survival (**c**) of SLR14- and vehicle-treated K18-hACE2 mice from 1 to 14 DPI. **d,e**, Measurement of genomic viral RNA in the lung 5 DPI by reverse-transcription quantitative PCR (RT-qPCR) against SARS-CoV-2 N gene using CDCN1 (**d**) or CDCN2 (**e**) primer-probe sets. **f**, Measurement of infectious virus titer in the lung 5 DPI by plaque assay. Limit of detection (LOD): 10^2^ PFU/mL. **g-i**, Measurement of expression of interferon stimulated genes (ISG) *Cxcl9* (**g**), *Isg15* (**h**) and *Usp18* (**i**) in the lung 5 DPI by RT-qPCR. **j,k**, Frequency of CD11b^+^CD64^+^ macrophages of CD45^+^ cells (**j**) and mean fluorescence intensity of MHCII on Ly6C^high^ monocytes (**k**) in the lung 5 DPI by flow cytometry. Mean ± s.e.m.; Statistical significance was calculated by log-rank Mantel–Cox test (**c**) or one-way ANOVA followed by Tukey correction (**d-k**); *P ≤ 0.05, **P ≤ 0.01, ***P ≤ 0.001, ****P ≤ 0.0001. Data are representative of two independent experiments.

To investigate the mechanisms by which SLR14 mediates protection, we collected lung tissues from naïve as well as infected mice treated with SLR14 or vehicle 5 DPI. Given the crucial role for RIG-I activation in the initiation of antiviral immunity, we first assessed the impact of SLR14 treatment on lung viral burden. We observed striking reduction in the level of genomic RNA (by RT-qPCR) and complete clearance of infectious virus (by plaque assay) in lung tissues from SLR14-treated mice compared to vehicle control (**Fig. 1d-f**). These results confirm that SLR14 affords protection against SARS-CoV-2 by efficiently mediating viral clearance in the lung tissue. Consistent with the absence of infectious virus, we found significantly attenuated expression of ISGs, including *Cxcl9*, *Isg15*, and *Usp18*, in lung tissues from SLR14-treated mice at this timepoint (**Fig. 1g-i**). In contrast, abundant ISG expression was detected in lungs from vehicle-treated mice, likely resulting from high viral burden.

To further probe the impact of SLR14 during SARS-CoV-2 infection, we assessed the immune infiltrates in lung tissues from these mice. We observed markedly decreased CD11b^+^Ly6C^+^ monocyte-derived proinflammatory macrophages in SLR14-treated mice 5 DPI (**Fig. 1j**). Additionally, SLR14 treatment led to a significant reduction in the surface expression of MHC class II molecules on Ly6C^high^ monocytes (**Fig. 1k**). Together, these results indicated that in addition to providing viral control, SLR14 alleviates lung immunopathology during SARS-CoV-2 infection by preventing viral replication.

## SLR14-mediated protection against SARS-CoV-2 depends on IFN-I signaling

To determine the molecular pathway required for SLR14-mediated protection against SARS-CoV-2, we investigated the effect of IFN-I signaling blockade using neutralizing antibodies against the receptor for IFN-I, interferon-α/β receptor (IFNAR) (**Fig. 2a**). Similar to SLR14 treatment at 4 hours post infection, K18-hACE2 mice were completely protected from morbidity and mortality when treated with SLR14 2 hours prior to SARS-CoV-2 infection. However, mice that were additionally pre-treated with anti-IFNAR antibodies lost the protection provided by SLR14 and all succumbed to the infection by 8 DPI (**Fig. 2b-d**). These results indicated that SLR14-mediated disease protection depends on IFN-I signaling.

**Figure 2.**
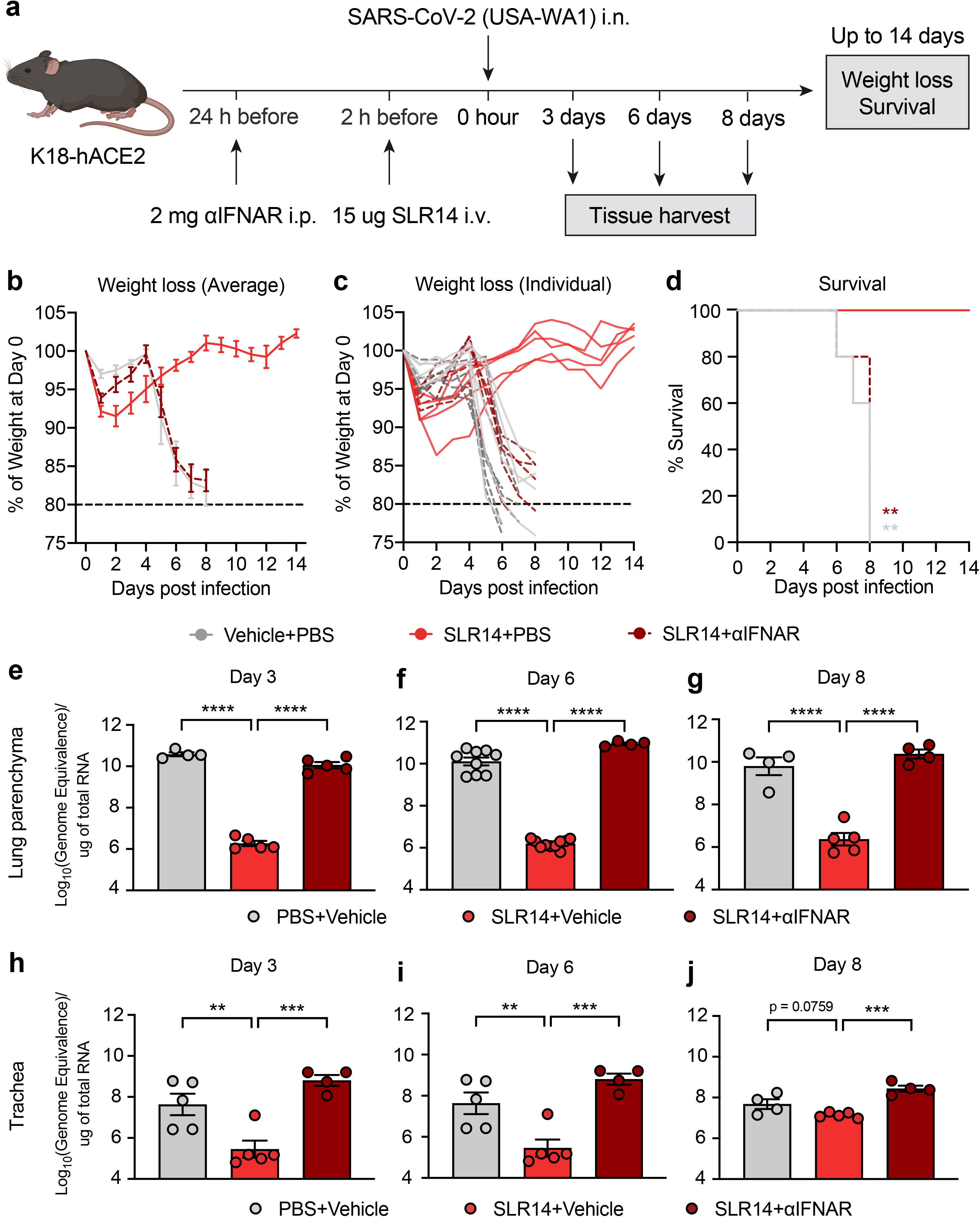
**a**, Experimental scheme: K18-hACE2 mice were intranasally infected with 10^3^ PFU SARS-CoV-2 (2019n-CoV/USA_WA1/2020). 2 hours before infection, 15 μg SLR14 or vehicle were intravenously administered. 24 hours before SLR14 injection, half of SLR14-treated mice was additionally given 2 mg anti-IFNAR antibodies. Weight loss and survival were monitored daily up to 14 DPI. Death was recorded when mice were found dead in the cage, moribund, or at 80% of original body weight. In a separate cohort, lung tissues were collected for virological analysis 3, 6, and 8 DPI. **b-d**, Weight loss (**b,c**) and survival (**d**) of SLR14- and vehicle-treated K18-hACE2 mice from 1 to 14 DPI. **e-g**, Measurement of genomic viral RNA in the lung parenchyma 3, 6, and 8 DPI by RT-qPCR using the CDCN2 primer-probe set. **h-j**, Measurement of genomic viral RNA in the trachea 3, 6, and 8 DPI by RT-qPCR using the CDCN2 primer-probe set. Mean ± s.e.m.; Statistical significance was calculated by log-rank Mantel–Cox test (**d**) or one-way ANOVA followed by Tukey correction (**e-j**); *P ≤ 0.05, **P ≤ 0.01, ***P ≤ 0.001, ****P ≤ 0.0001. Data are representative of two independent experiments.

Viral infections of the lower respiratory tract are a leading cause of mortality in this disease context, whereas upper respiratory infection primarily contributes to viral transmission. To characterize the tissue sites that are protected by SLR14 and the contribution of IFN-I signaling to SLR14-mediated protection, we collected the lung parenchyma, the trachea, and a nasal wash to assess viral burden throughout the respiratory tract at 3, 6, and 8 DPI. The ability of SLR14 to suppress lung viral replication was prominent as early as 3 DPI and maintained throughout the course of infection up to 8 DPI. However, the reduction in the level of viral genomic RNA in lung tissues was completely abolished when mice were also pre-treated with anti-IFNAR antibodies (**Fig. 2e-g**). These findings were largely recapitulated in the trachea, although the overall viral titer was lower than that of the lung (**Fig. 2h-j**). We also observed a significant decrease in the level of genomic RNA in nasal washes from SLR14-treated mice at 8 DPI, although not at 3 or 6 DPI (**Extended Data Fig. 1a-d**). The lack of an apparent early antiviral effect in the nasal cavity highlights the spatial compartmentalization of the respiratory mucosa and suggests intranasal SLR14 application as an alternative route of delivery to achieve optimal protection in the upper respiratory tract. These results indicated that SLR14 utilizes IFN-I signaling to mediate viral clearance in the lower respiratory tract, thereby preventing severe respiratory disease.

We additionally assessed the role of IFN-I signaling in SLR14-mediated viral control using IFNAR-deficient (*Ifnar*^−/−^) mice. Laboratory mice are not susceptible to SARS-CoV-2 infection due to the inability of the virus to utilize the mouse orthologue of human ACE2 for viral entry. Therefore, we first transduced C57BL/6J (B6J) or *Ifnar*^−/−^ mice with hACE2-expressing adeno-associated viruses (AAV-hACE2) through intratracheal delivery to sensitize them for SARS-CoV-2 infection (**Extended Data Fig. 2a**) ^34^. Two weeks post transduction, AAV-hACE2 B6J or *Ifnar*^−/−^ mice were infected with SARS-CoV-2 and treated with SLR14 4 hours post infection. Consistent with experiments using anti-IFNAR antibodies, AAV-hACE2 *Ifnar*^−/−^ mice did not respond to SLR14 and maintained high levels of viral genomic RNA similar to that of untreated controls 4 DPI (**Extended Data Fig. 2b**). In contrast, SLR14-treated AAV-hACE2 B6J mice had significantly reduced level of genomic RNA compared to untreated controls. Together, these results showed, by two separate approaches, that the SLR14-mediated antiviral resistance against SARS-CoV-2 requires IFNAR.

## SLR14 is taken up by various cell types in the lung

RIG-I is ubiquitously expressed in all cell types^35^. To determine the cell type that is being targeted by PEI-complexed SLR14 following i.v. injection and responsible for producing an early source of IFN-I to mediate protection, we injected Alexa Flour 647 (AF647)-conjugated SLR14 into naïve K18-hACE2 mice, and collected lung tissues 4 hours post injection to assess cellular uptake of SLR14 by flow cytometry (**Extended Data Fig. 3a**). Of the total SLR14^+^ cells, we found SLR14 to be broadly distributed across multiple immune and non-immune cellular compartments (**Extended Data Fig. 3b**). In particular, EpCAM^+^ epithelial cells and CD64^+^ macrophages accounted for the majority of SLR14 uptake (∼70% of SLR14^+^ cells). We further analyzed the composition of SLR14^+^ macrophages and found this population to be mainly CD11b^+^Ly6C^+^ monocyte-derived proinflammatory macrophages, although some SLR14^+^ interstitial and alveolar macrophages were also found (**Extended Data Fig. 3c**). We additionally derived a distribution index to account for cell type abundance and observed similar patterns of SLR14 uptake by epithelial cells and macrophages (**Extended Data Fig. 3d-e**). Together, these results indicated that i.v. injected SLR14 is mainly taken up by lung epithelial cells and inflammatory macrophages, contributing to the rapid production of IFN-I and elicitation of local ISG response against SARS-CoV-2 infection.

## SLR14 treatment timing relative to SARS-CoV-2 infection determines protective activities

Early and robust IFN-I production in response to infection with SARS-CoV-2 is essential for rapid control of viral replication, whereas IFN-I induced late during the infection may contribute to immunopathology and drive severe disease. Thus, we next examined the effect of treatment timing on the protective capacity of SLR14. We treated K18-hACE2 mice with SLR14 at different timepoints relative to SARS-CoV-2 challenge (**Fig. 3a**). Prophylactic treatment of SLR14 either at 16 or 2 hours before infection protected mice from weight loss and clinical disease after SARS-CoV-2 infection (**Fig. 3b-d**). Similarly, treatment of SLR14 4 hours after infection as a post-exposure prophylaxis was also highly protective and largely prevented disease development. However, the efficacy of SLR14 became more dependent on treatment timing when administered therapeutically. Treatment at 24 or 48 hours post infection resulted in an intermediate level of protection (40% survival) with some level of morbidity and mortality being observed, while SLR14 lost its protective capacity when administered 72 hours post infection (**Fig. 3e-g**). These results corroborated the protective role of early IFN-I, and importantly, demonstrated that SLR14-based treatment can be broadly used as prophylaxis and early post-exposure prophylaxis against COVID-19.

**Figure 3.**
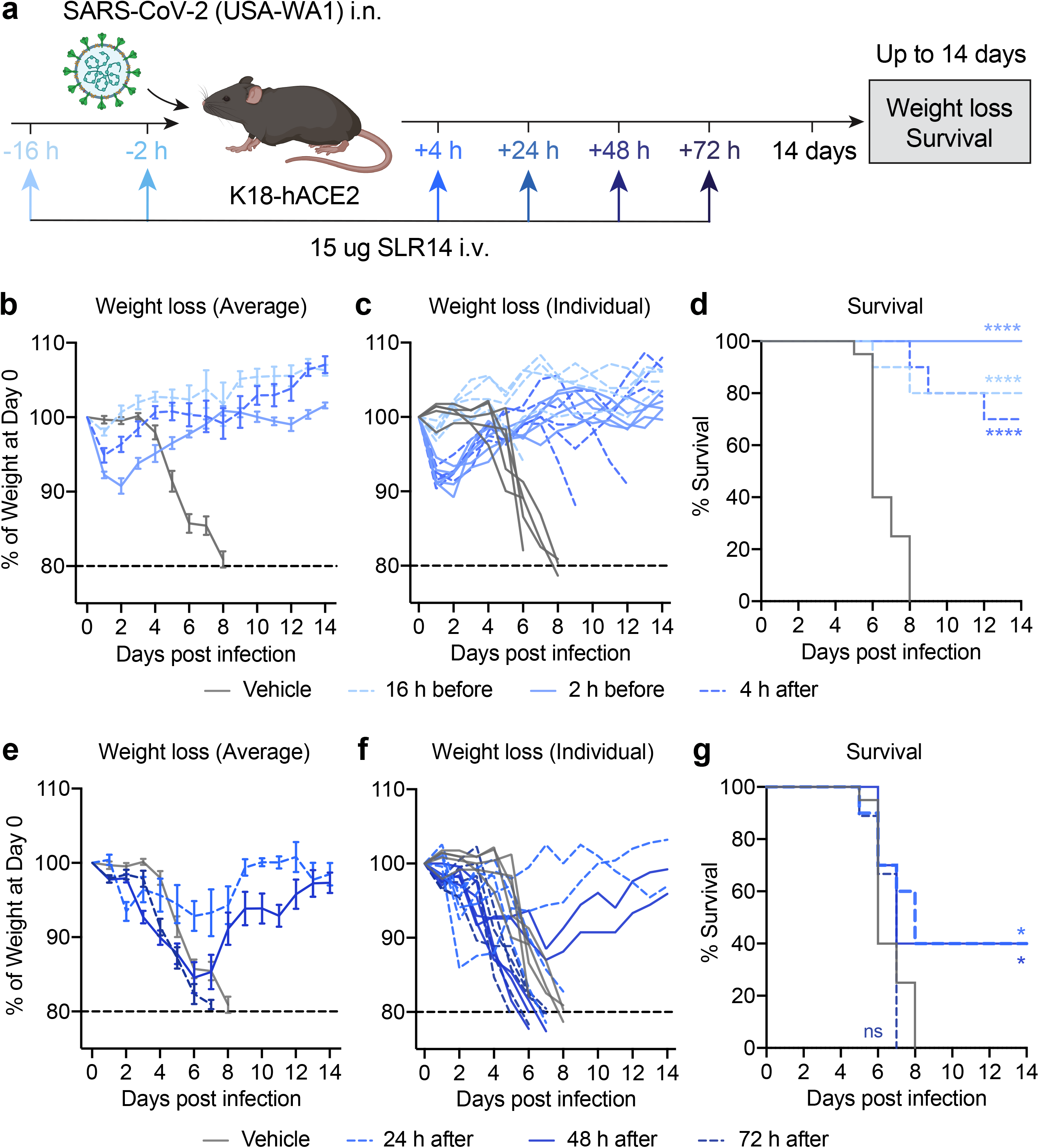
**a**, Experimental scheme: K18-hACE2 mice were intranasally infected with 10^3^ PFU SARS-CoV-2 (2019n-CoV/USA_WA1/2020). 15 μ SLR14 were intravenously administered at 16 hours before, 2 hours before, 4 hours after, 24 hours after, 48 hours after, or 72 hours after infection. Weight loss and survival were monitored daily up to 14 DPI. Death was recorded when mice were found dead in the cage, moribund, or at 80% of original body weight. **b-d**, Weight loss (**b,c**) and survival (**d**) of prophylactically SLR14- and vehicle-treated K18-hACE2 mice from 1 to 14 DPI. **e-g**, Weight loss (**e,f**) and survival (**g**) of therapeutically SLR14- and vehicle-treated K18-hACE2 mice from 1 to 14 DPI. Mean ± s.e.m.; Statistical significance was calculated by log-rank Mantel– Cox test (**d,g**); *P ≤ 0.05, **P ≤ 0.01, ***P ≤ 0.001, ****P ≤ 0.0001. Data are pooled from of three independent experiments.

## Therapeutic SLR14 cures persistent SARS-CoV-2 infection in immunodeficient mice through induction of IFN-I

There is a clinically unmet need for the development of an effective therapy to treat chronic SARS-CoV-2 infection in immunodeficient individuals and prevent further emergence of viral variants. We have previously demonstrated that AAV-hACE2-transduced *Rag1*^−/−^ or *Rag2*^−/−^ mice (which completely lack mature T and B cells, collectively referred to as *Rag*^−/−^ mice) become chronically infected following SARS-CoV-2 infection, similar to what is seen in immunodeficient patients^36^. These mice maintain stable levels of viral RNA and infectious virus for at least 14 DPI. This is in stark contrast to B6J mice, which clear the infection by 7 DPI and remain virus free thereafter. Given that convalescent plasma (CP) therapy has been implemented to treat immunocompromised patients with COVID-19^37^, we first validated whether persistently infected *Rag*^−/−^ mice are a clinically relevant model in their response to CP therapy. To this end, we adoptively transferred sera from convalescent AAV-hACE2 B6J mice into persistently infected recipient AAV-hACE2 *Rag*^−/−^ mice 7 DPI and measured lung viral titer 14 DPI (**Extended Data Fig. 4a**). We found that CP transfer resulted in significant reduction in genomic RNA and complete clearance of infectious virus in the lung, compared with PBS-treated SARS-CoV-2 infected AAV-hACE2 *Rag*^−/−^ controls (**Extended Data Fig. 4b,c**). These results suggest that *Rag*^−/−^ mice is a suitable *in vivo* model of immunocompromised patients for preclinical testing of antiviral therapeutics, as they support persistent SARS-CoV-2 infection and derive benefits from CP therapy.

We next examined whether SLR14 can be used as a therapeutic modality to treat persistent infection in *Rag*^−/−^ mice. We infected AAV-hACE2 *Rag*^−/−^ mice with SARS-CoV-2, treated them with SLR14 7 DPI, and collected lung tissues 14 DPI to assess the viral burden (**Fig. 4a**). 1 day prior to SLR14 treatment, a subset of SLR14-treated AAV-hACE2 mice also received anti-IFNAR antibodies. SLR14 treatment led to a significant reduction in the level of genomic viral RNA in the lung (**Fig. 4b**). The ability of SLR14 in decreasing viral RNA was abolished when anti-IFNAR blocking antibody was given, suggesting SLR14 similarly utilizes IFN-I signaling to promote viral clearance in mice lacking the adaptive immune system. We additionally observed a striking difference in the infectious viral load in lung tissues from SLR14-treated mice compared to vehicle controls. Treatment with SLR14, but not vehicle, significantly reduced viral burden and resulted in complete clearance of infectious virus in 5 out of 7 AAV-hACE2 *Rag*^−/−^ mice and reduction of viral titer in the remaining 2 (**Fig. 4c**). Moreover, SLR14-mediated protection required IFNAR signaling (**Fig. 4c**). These results showed that in the setting of complete T- and B-cell-deficiency, a single therapeutic SLR14 treatment, through the induction of IFN-I, is sufficient to cure persistent infection.

**Figure 4.**
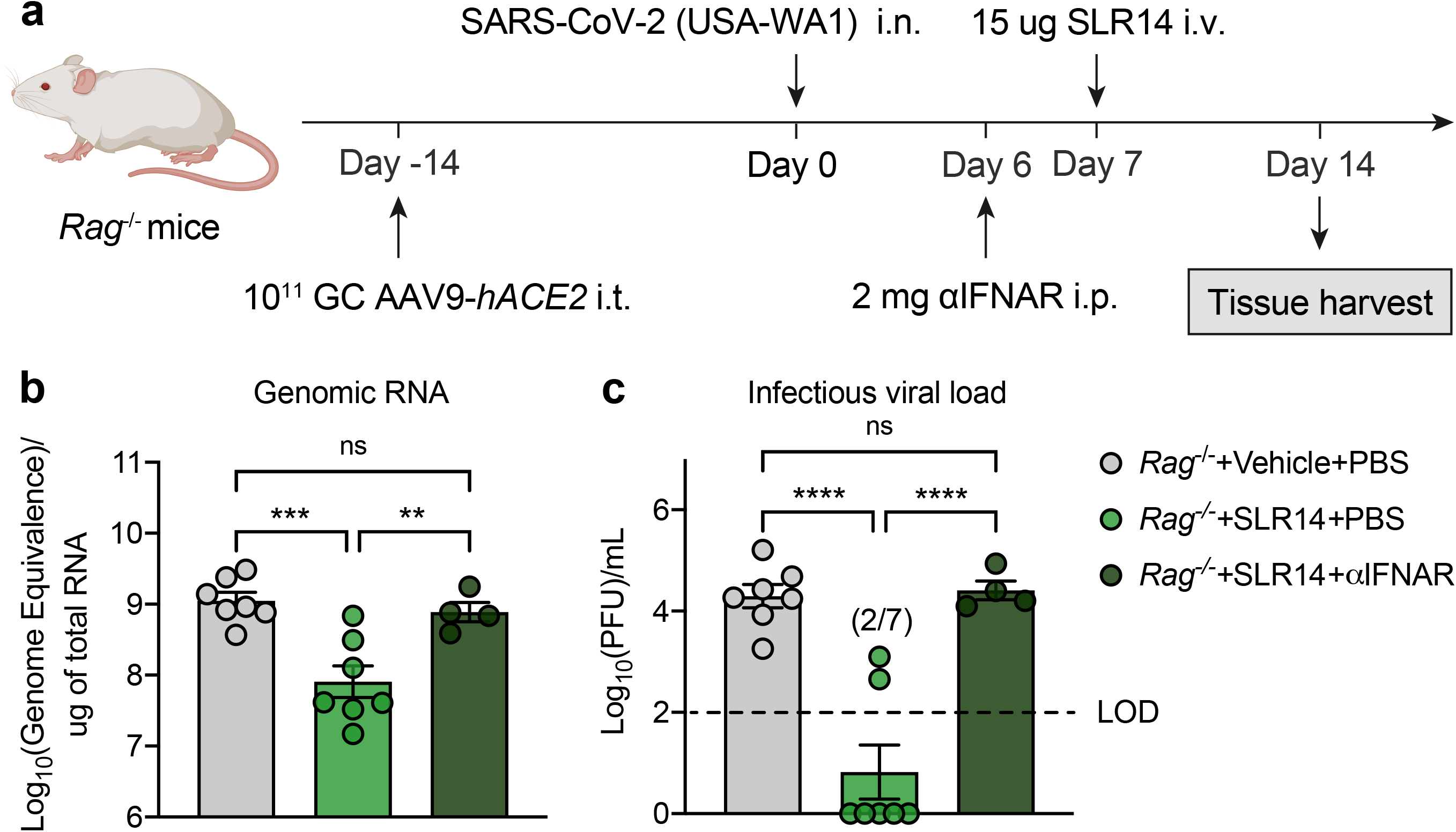
**a**, Experimental scheme: *Rag*^−/−^ mice were intratracheally administered with 10^11^ genome copies of AAV9-*hACE2* and let rest for 2 weeks before intranasal infection with 10^6^ PFU SARS-CoV-2 (2019n-CoV/USA_WA1/2020). 15 μg SLR14 or vehicle were intravenously administered 7 DPI. 24 hours before SLR14 injection, half of SLR14-treated mice was additionally given 2 mg anti-IFNAR antibodies. Lung tissues were collected for virological analysis 14 DPI. **b**, Measurement of genomic viral RNA in the lung 14 DPI by RT-qPCR. **c**, Measurement of infectious virus in the lung 14 DPI by plaque assay. Limit of detection (LOD): 10^2^ PFU/mL. Mean ± s.e.m.; Statistical significance was calculated by one-way ANOVA followed by Tukey correction (**b,c**); *P ≤ 0.05, **P ≤ 0.01, ***P ≤ 0.001, ****P ≤ 0.0001. Data are pooled from two independent experiments.

## SLR14 affords broad protection against immunologically evasive SARS-CoV-2 variants

As SARS-CoV-2 variants continue to emerge and spread, antiviral therapeutics that confer broadly cross-reactive protection are urgently needed. Emerging evidence suggests that several SARS-CoV-2 variants have acquired mutations that confer elevated resistance to IFN-I treatment in cell culture^38^. However, whether such altered properties *in vitro* translate into evasion of IFN-based therapy *in vivo* remains unclear. To this end, we obtained 4 clinically relevant SARS-CoV-2 VOC, including P.1, B.1.526, B.1.1.7, and B.1.351, and used them to infect K18-hACE2 mice. The P.1, B.1.526, and B.1.1.7 variants were identified and isolated as part of the Yale SARS-CoV-2 Genomic Surveillance Initiatives^39^, and the B.1.351 variant was obtained from BEI Resources Repository. All variants were confirmed to harbor signature mutations characteristic of their respective lineages and show correct placement in the phylogenetic tree built with public SARS-CoV-2 genomic sequences (**Extended Data Table 1, Extended Data Fig. 5**). To examine whether SLR14 is protective against SARS-CoV-2 VOC, we infected mice with variants P.1, B.1.351, B.1.526, or B.1.1.7, and treated them with SLR14 4 hours post infection (**Fig. 5a**). SLR14 afforded potent protection against the P.1 variant, almost completely preventing morbidity and mortality in the face of highly pathogenic infection, even when treated post-exposure prophylactically (**Fig. 5b-d**). Similarly, SLR14 fully prevented weight loss or any discernable disease following infection with B.1.526, which, in untreated mice, caused a less pathogenic infection compared to that of the ancestral strain or circulating variants (**Fig. 5e-g**). Consistent with the reported resistance to IFN-I signaling *in vitro*^38^, SLR14 treatment was less effective against infection with B.1.351 or B.1.1.7 *in vivo*, conferring approximately 40 to 50% net protection in K18-hACE2 mice (60% survival in SLR14-treated mice over 10 to 20% survival in vehicle controls) (**Fig. 5h-m**). Additional experiments with the B.1.1.7 variant confirmed its partial resistance to SLR14 treatment, irrespective of initial sizes of viral inoculum (**Extended Data Fig. 6a-f**). Nevertheless, clear benefits were seen with prophylactic treatment of SLR14. These results suggest that SLR14 confers broad-coverage protection against antibody- and IFN-I-evasive variants.

**Figure 5.**
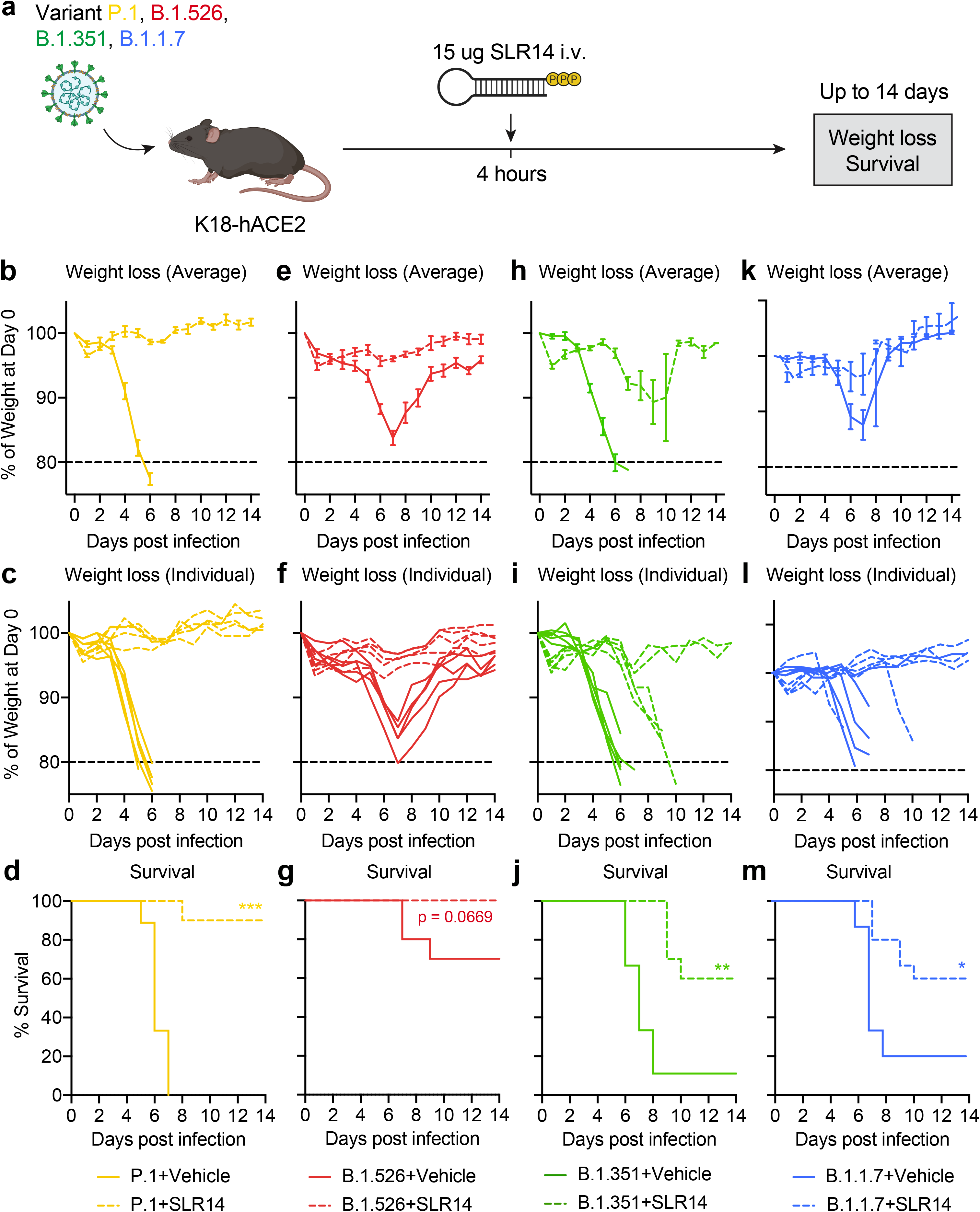
**a**, Experimental scheme: K18-hACE2 mice were intranasally infected with 10^4^ PFU P.1, B.1.526, B.1.351, or B.1.1.7 variants. 15 μ SLR14 or vehicle were intravenously administered at 4 hours after infection. Weight loss and survival were monitored daily up to 14 DPI. Death was recorded when mice were found dead in the cage, moribund, or at 80% of original body weight. **b-d**, Weight loss (**b,c**) and survival (**d**) of SLR14- and vehicle-treated K18-hACE2 mice from 1 to 14 DPI following P.1 infection. **e-g**, Weight loss (**e,f**) and survival (**g**) of SLR14- and vehicle-treated K18-hACE2 mice from 1 to 14 DPI following B.1.526 infection. **h-j**, Weight loss (**h,i**) and survival (**j**) of SLR14- and vehicle-treated K18-hACE2 mice from 1 to 14 DPI following B.1.351 infection. **k-m**, Weight loss (**k,l**) and survival (**m**) of SLR14- and vehicle-treated K18-hACE2 mice from 1 to 14 DPI following B.1.1.7 infection. Mean ± s.e.m.; Statistical significance was calculated by log-rank Mantel–Cox test (**d,g,j,m**); *P ≤ 0.05, **P ≤ 0.01, ***P ≤ 0.001, ****P ≤ 0.0001. Data are pooled from two independent experiments.

## Discussion

The sudden arrival and devastating spread of COVID-19 have emphasized the importance of continuous efforts to develop broad-spectrum antiviral agents. Here, we examined the *in vivo* efficacy of SLR14 against viral replication throughout the respiratory tract and disease development in a mouse model of severe SARS-CoV-2 infection. We found that SLR14 conferred considerable antiviral resistance in the lower respiratory tract and effectively prevented morbidity and mortality following infection with the ancestral virus. We also examined the effect of host factor, tissue compartment, and treatment timing in the protective capacity of SLR14, and found that the protective efficacy of SLR14 depends on intact IFNAR signaling and showed early SLR14 administration provided superior protection, while treatment as late as 48 hours post infection still afforded partial protection. We further tested the therapeutic potential for SLR14 in chronically infected immunodeficient mice and demonstrated that a single dose of SLR14 conferred near-sterilizing immunity by the innate immune system alone, even in the absence of T and B cells. Finally, we found that SLR14 confers broad protection against all emerging SARS-CoV-2 variants.

The apparent protective role of early and regulated IFN-I suggests IFN-based therapies can be utilized for prevention and treatment of COVID-19. In a golden hamster model of SARS-CoV-2 infection, intranasal administration of commercially available universal IFN (IFNα A/D) reduced viral burden and attenuated pathology in the lung40. Growing evidence also suggests that recombinant IFN-based intervention during the early stage of COVID-19 could provide desired clinical benefits in humans. In a retrospective multicenter cohort study of 446 COVID-19 patients, early administration of inhaled IFNα2b produced more favorable clinical responses compared to lopinavir/ritonavir (LPV/r) treatment alone and was associated with reduced in-hospital mortality41. While results from recombinant IFN-based clinical trials are promising, one of the major disadvantages of this approach is its high cost, with the direct medical cost of IFN treatment regimen to range between $1,120 and $1,962 and PEG-IFN treatment regimen between $2,156 and $5,88742. In addition, administration of recombinant IFN has been shown to induce neutralizing antibody response that could render the therapy ineffective43. SLR14 addresses these challenges with 1) drastically increased affordability due to its synthetic simplicity, small size, and manufacturability, and 2) its ability to induce different members of the IFN-I family, including 10 IFN-α subtypes and an IFN-β, which maximizes the likelihood of downstream responses to be functional19. In particular, the ability of SLR14 to elicit IFN-β is ideal as it enables early medical intervention for COVID-19 patients with pre-existing autoantibodies against one or multiple subtypes of IFN-α, who are particularly susceptible to prolonged viral replication and severe disease after infection with SARS-CoV-244.

The clinical efficacy of CP in patients with severe COVID-19 has not been thoroughly demonstrated, and its use in different stages of infection and disease remains experimental^45,46^. Emergence of immune-evading variants from patients with immunosuppression of T cell and B cell arms indicate caution should be used for CP therapy^47^. In these patients, the administered antibodies have little support from cytotoxic CD8 T cells or helper CD4 T cells, thereby reducing the chances of clearance and theoretically allowing for SARS-CoV-2 escape. Therefore, a novel therapeutic paradigm that treats persistent viral infection regardless of its effect on adaptive immunity will hold immense potential for this patient population. We have shown in this study that a single dose of SLR14 in mice lacking the adaptive immune system in the setting of chronic SARS-CoV-2 infection can induce near-sterilizing immunity. These results demonstrated that SLR14’s utility extends beyond prophylactic antivirals, but also therapeutics that can be given to patients with immunocompromised conditions, providing an immediate solution to simultaneously cure chronic infection and suppress future emergence of immune-evasive variants. From an evolutionary perspective, such sterilizing protection induced exclusively through innate immune activation is analogous to antiviral mechanisms in metazoan organisms lacking adaptive immunity, which provides a basic, yet crucial, protective strategy against viral pathogens^48^.

Vaccines remain the best approach to thwart the COVID-19 pandemic. However, with many countries lacking access to adequate vaccine doses, alternative strategies need to be developed and rapidly distributed to parts of the world severely impacted by these variants. Here, we showed that SLR14 potently prevented morbidity and mortality following infection with clinically relevant VOC, which have vastly different signature mutations and immune-evading capacity. Consistent with a recent study examining IFN-I potency against different variants *in vitro*^38^, the protective capacity of SLR14 was less impressive when administered against B.1.351 or B.1.1.7. Importantly, SLR14 still retained considerable residual antiviral capacity, which may be attributed by the speed, magnitude, and diversity of IFN-I responses induced by SLR14 that could collectively overcome viral resistance^19^. Up until this point, B.1.1.7 has been recognized as a minimally immune-evasive variant based on both antibodies and T cell recognition. Here, we provide the first set of *in vivo* evidence to suggest that B.1.1.7 exhibits signs of IFN-I evasion and responds only moderately to IFN-based therapy. Such innate immune evasion may underlie the rapid global spread of B.1.1.7. These results showcase SLR14’s ability to not only be utilized as a therapeutic agent, but also an investigative tool for functional assessment of basic SARS-CoV-2 biology.

Several drugs have been approved by FDA under Emergency Use Authorization (EUA), including dexamethasone, remdesivir, monoclonal antibodies, tocilizumab, and baricitinib, to treat COVID-19^3,4, 49–52^. However, these therapeutics typically provide modest benefits at best and are limited to a subset of patients. While currently licensed vaccines demonstrate astounding protective efficacy against COVID-19, a new variant may develop in the future to significantly reduce efficacy. Further, there is a global shortage in vaccines with inequitable access in many lower income countries. The development, characterization, and ultimate deployment of an effective antiviral against SARS-CoV-2 could prevent substantial morbidity and mortality associated with COVID-19. In addition to its therapeutic potentials, SLR14 can be used as an invaluable investigative tool to advance our understanding of protective antiviral immunity against respiratory viruses, which will enable the rational design of next-generation antiviral therapeutics. Lastly, with the prevalence of prepandemic zoonotic viruses and the unpredictable overlap of human and wild animal ecologies, the potential for a novel viral emergence from its natural reservoir into humans is a matter of time. Through the creation of a simple and versatile RNA-based therapeutics, our studies will facilitate pandemic preparedness and response against future respiratory pathogens.

## Materials and Methods

### Ethics

The Institutional Review Board from the Yale University Human Research Protection Program determined that the RT-qPCR testing and sequencing of de-identified remnant COVID-19 clinical samples conducted in this study is not research involving human subjects (IRB Protocol ID: 2000028599).

### Mice

B6.Cg-Tg(K18-ACE2)2Prlmn/J (K18-hACE2), B6(Cg)-*Ifnar1*^tm1.2Ees^/J (*Ifnar1*^−/−^), B6.129S7-*Rag1^tm1Mom^*/J (B6J *Rag1*^−/−^), and C.129S7(B6)-*Rag1^tm1Mom^*/J (BALB/c *Rag1*^−/−^) mice were purchased from the Jackson Laboratories and were subsequently bred and housed at Yale University. *Rag2*^−/−^ mice were generously gifted from R. Flavell (Yale University). 6- to 10-week-old mixed sex mice were used throughout the study. All mice were housed as groups of 5 to 6 individuals per cage and maintained on a 12-hour light/dark cycle (lights on at 7:00 AM) at 22–25°C temperature and 30–70% relative humidity under specific-pathogen free conditions. All mice were fed with regular rodent’s chow and sterilized water ad libitum. All procedures used in this study (sex-matched, age-matched) complied with federal guidelines and the institutional policies of the Yale School of Medicine Animal Care and Use Committee.

### Virus sequencing

Nucleic acid was extracted from 300 µL viral transport medium from nasopharyngeal swabs and eluted in 75 µL using the MagMAX viral/pathogen nucleic acid isolation kit. Extracted nucleic acid was tested by our multiplexed RT-qPCR variant assay^53,54^, and then libraries were prepared using the Illumina COVIDSeq Test RUO version. The protocol was slightly modified by lowering the annealing temperature of the amplicon PCR step to 63°C, and by reducing tagmentation to 3 minutes. Pooled libraries were sequenced on the Illumina NovaSeq (paired-end 150). Data was processed and consensus sequences were generated using iVar (version 1.3.1) with the minimum depth threshold (-m) at 20 and minimum frequency threshold (-t) at 0.6^55^. Genome sequences were uploaded to GISAID. Samples belonging to the B.1.1.7 (EPI_ISL_1038987), P.1 (EPI_ISL_1293215), and B.1.526 (EPI_ISL_944591) lineages were selected for virus isolation from the original sample. Virus belonging to the B.1.351 lineage was obtained from BEI Resources.

### Virus isolation

Samples selected for virus isolation were diluted 1:10 in Dulbecco’s Modified Eagle Medium (DMEM) and then filtered through a 45 µM filter. The samples were ten-fold serially diluted from 1:50 to 1:19,531,250. The dilution was subsequently incubated with TMPRSS2-Vero E6 in a 96 well plate and adsorbed for 1 hour at 37°C. After adsorption, replacement medium was added, and cells were incubated at 37°C for up to 5 days. Supernatants from cell cultures with cytopathic effect (CPE) were collected, frozen, thawed and subjected to RT-qPCR. Fresh cultures were inoculated with the lysates as described above for viral expansion. Viral infection was subsequently confirmed through reduction of Ct values in the cell cultures with the multiplex variant qPCR assay. Expanded viruses were re-sequenced following the same method as described above and were identical to the original clinical sample sequence. Genome sequences of cultured viruses B.1.1.7 (SARS-CoV-2/human/USA/Yale-3363/2021; GenBank accession: MZ202178), B.1.351 (SARS-CoV-2/human/ZAF/Yale-3366/2020; GenBank accession: MZ202314), P.1 (SARS-CoV-2/human/USA/Yale-3365/2021; GenBank accession: MZ202306), and B.1.526 (SARS-CoV-2/human/USA/Yale-3362/2021; GenBank accession: MZ201303) were uploaded to GenBank. We used Nextclade v0.14.2 (https://clades.nextstrain.org/) to generate a phylogenetic tree and to compile a list of amino acid changes in the virus isolates as compared to the Wuhan-Hu-1 reference strain (**Extended Data Table 1; Extended Data Fig. 5**). Lineage defining amino acid changes were marked based on the outbreak.info lineage comparison^56^.

### Synthesis, purification, and labeling of the SLR-14 oligonucleotide

The triphosphorylated RNA oligonucleotides SLR-14 (5′ pppGGAUCGAUCGAUCGUUCGCGAUCGAUCGAUCC-3′ and SLR-14-amino (5′ pppGGAUCGAUCGAUCGUXCGCGAUCGAUCGAUCC-3′ where X = aminomodifier C6dT; Glen Research) were prepared as described^57^. Briefly, for every 1 mg of starting material, removal of the oligonucleotide from the polymer support and base deprotection was performed in a 1:1 mixture of 40% methylamine (Sigma-Aldrich) and 30% ammonium hydroxide (JT Baker) at 65°C for 15 min. The solution was cooled on ice for 10 min, transferred to a new vial, and evaporated to dryness. 500 µL of absolute ethanol was added, and the mixture was evaporated to dryness again. To deprotect the 2′-OH groups, the dry oligonucleotide was incubated with 500 µL of a 1 M solution of tetrabutylammonium fluoride in tetrahydrofuran (Sigma-Aldrich) at room temperature for 36 h. 500 µL of 2 M sodium acetate (pH 6.0) was added, and the solution was evaporated to a 500–600 µL volume, extracted with 3 × 800 µL of ethyl acetate, and ethanol precipitated. The RNA oligonucleotide was then purified on a 16% denaturing polyacrylamide gel. For fluorescent labeling, for every 1 mg of starting material, the purified SLR14-amino oligonucleotide was dissolved in 200 µL of 0.25 M sodium bicarbonate buffer (pH 9.2). Then, a solution containing 0.5 mg of Alexa Fluor 647 NHS ester (Life Technologies Corp.) in 200 µL N,N-dimethylformamide was added, and the reaction mixture was incubated at room temperature for 2 h. The labeled oligonucleotide (AF647-SLR14) was ethanol precipitated and purified on a 20% denaturing polyacrylamide gel.

### SARS-CoV-2 mouse infection and antibody treatment

Before infection, mice were anesthetized using 30% (vol/vol) isoflurane diluted in propylene glycol. For K18-hACE2 mice, 50 µL of SARS-CoV-2 was delivered intranasally at 10^3^ PFU per mouse (LD_100_), unless specified otherwise. Following infection, weight loss and survival were monitored daily up to 14 DPI. For AAV-hACE2 mice, 50 µL of SARS-CoV-2 was delivered intranasally at 10^6^ PFU per mouse. For IFNAR blockade, mice were treated once with 2 mg of blocking antibodies one day prior to infection (Clone MAR1-5A3; BioXCell). Experiments involving SARS-CoV-2 infection were performed in a Biosafety Level 3 (BSL-3) facility with approval from the Yale Institutional Animal Care and Use Committee (IACUC) and Yale Environmental Health and Safety (EHS).

### Intravenous injection of SLR14 in mice

At indicated timepoints, 15 µg SLR14 was i.v. injected. Briefly, 15 µg (∼0.6 mg/kg body weight) SLR14 and 4 µL jetPEI (Polyplus Transfection) were diluted and mixed with 5% glucose solution to a total of 100 µL injection solution per mouse. After 15 min of incubation at room temperature, the 100-µL complex was carefully injected into the retro-orbital sinus with a 0.5-mL BD Insulin syringe. Before injection, mice were anesthetized using 30% (vol/vol) isoflurane diluted in propylene glycol. H_2_O and jetPEI were mixed with 5% glucose solution and used as a vehicle control.

### AAV-hACE2 transduction

AAV9 vector encoding hACE2 was purchased from Vector Biolabs (AAV9-CMV-hACE2). Animals were anaesthetized using a mixture of ketamine (50 mg/kg) and xylazine (5 mg/kg), injected intraperitoneally. The rostral neck was shaved and disinfected. A 5-mm incision was made, the salivary glands were retracted, and the trachea was visualized. Using a 32-G insulin syringe, a 50-µL bolus injection of 10^11^ genomic copies AAV-CMV-hACE2 was injected into the trachea. The incision was closed with 4–0 Vicryl suture. Following intramuscular administration of analgesic (meloxicam and buprenorphine, 1 mg/kg), animals were placed on a heating pad and closely monitored until full recovery.

### Adoptive transfer of sera

WT AAV-hACE2 mice were infected with SARS-CoV-2 as indicated above. At 14 DPI animals were euthanized for blood collection. Blood was allowed to coagulate at room temperature for 30 min and then was centrifuged at 3900 rpm for 20 min at 4°C. Serum was collected, and anesthetized mice (30% v/v Isoflurane diluted in propylene glycol) were injected with 200 µL serum with a 32 g 8 mm syringe via retro orbital route.

### Measurements of genomic RNA and infectious virus

Viral RNA and titer from mouse lung tissues were measured as previously described^34^. Briefly, at indicated time points, mice were euthanized with 100% isoflurane. The whole lung was placed in a Lysing Matrix D tube (MP Biomedicals) with 1 mL of PBS, and homogenized using a table-top homogenizer at medium speed for 2 min. For RNA analysis, 250 μL of the lung homogenates was added to 750 μL Trizol LS (Invitrogen), and RNA was extracted with the RNeasy Mini Kit (Qiagen) according to the manufacturer’s instructions. SARS-CoV-2 RNA levels were quantified with 250 ng of RNA inputs using the Luna Universal Probe One-Step RT-qPCR Kit (New England Biolabs), using real-time RT-PCR primer/probe sets 2019-nCoV_N1 (CDCN1) and 2019-nCoV_N2 (CDCN2). For determination of infectious titer, plaque assays were performed using lung homogenates in Vero E6 cells cultured with MEM supplemented with NaHCO_3_, 4% FBS, and 0.6% Avicel RC-581. 48 h after infection, plaques were resolved by 1 h fixation with 10% formaldehyde and sequentially 1 h staining with 0.5% crystal violet in 20% ethanol. Finally, plates were rinsed in water for better visualization of plaques.

### Antibodies for flow cytometry

Anti-mouse antibodies used in this study, together with vendors and dilutions, are listed as follows: FITC anti-mCD11c (N418) (1:400) (BioLegend), PerCP-Cy5.5 anti-mLy6C (HK1.4) (1:400) (BioLegend), Alexa Fluor 700 anti-mLy6G (1A8) (1:400) (BioLegend), Brilliant Violet 786 anti-mCD11b (M1/70) (1:400) (BioLegend), APC-Cy7 anti-mTCRb (H57-597) (1:400) (BioLegend), APC-Cy7 anti-mCD3 (H57-597) (1:400) (BioLegend), APC-Cy7 anti-mCD19 (6D5) (1:400) (BioLegend), APC-Cy7 anti-mNK1.1 (PK136) (1:400) (BioLegend), PE anti-mCD64 (X54-5/7.1) (1:200) (BioLegend), Brilliant Violet 711 anti-mSiglecF (E50-2440) (1:200) (BD Biosciences), and Pacific Blue anti-mI-A/I-E (M5/114.15.2) (1:400) (BioLegend).

### Flow cytometry

Mouse lung tissues were collected at experimental end-point, digested with 1 mg/mL collagenase A (Roche) and 30 µg/mL DNase I (Sigma-Aldrich) in complete RPMI-1640 media for 30 min at 37 °C, and mechanically minced. Digested tissues were then passed through a 70 µm strainer (Fisher Scientific) to single cell suspension and treated with ACK Lysing Buffer (ThermoFisher) to remove red blood cells. Cells were resuspended in Live/Dead Fixable Aqua (ThermoFisher) for 20 min at 4 °C. Following L a wash, cells were blocked with anti-mouse CD16/32 antibodies (BioXCell) for 30 min at 4 °C. Cocktails of staining antibodies were added directly to this mixture for 30 min at 4 °C. Prior to analysis, mouse cells were washed and resuspended in 100 μL 4% PFA for 30–45 min at 4 °C. Following this incubation, cells were washed and prepared for L analysis on an Attune NXT (ThermoFisher). Data were analysed using FlowJo software version 10.6 software (Tree Star). The specific sets of markers used to identify each subset of cells are summarized in **Extended Data Fig. 7**.

### Statistical analysis

The data were analyzed by log-rank Mantel–Cox test or one-way ANOVA followed by Tukey correction. All statistical tests were calculated using GraphPad Prism (GraphPad software). A P value of <0.05 was considered statistically significant.

### Graphical illustrations

Graphical illustrations were made with Biorender.com.

## Acknowledgements

The authors gratefully acknowledge all members of the Iwasaki lab for invaluable feedback, especially Annsea Park for critical reading of the manuscript and Xiaodong Jiang for helpful discussions. We thank Barney Graham (NIH-VRC) for kindly providing VeroE6 cells overexpressing ACE2 and TMPRSS2. We give special recognition of the services of Ben Fontes and the Yale EHS Department for their on-going assistance in safely conducting BSL3 research. We also thank the Yale IMPACT Research Team, Yale-New Haven Hospital, and Connecticut Department of Public Health for their partnerships in our SARS-CoV-2 Genomic Surveillance Initiative. This work was supported by Women’s Health Research at Yale Pilot Project Program (A.I.), the Mathers Family Foundation (A.I., C.B.W.), the Ludwig Family Foundation (A.I., C.B.W.), NIAID 1R01AI157488-01 (A.I.), and NIAID 5R01AI127429-04 (A.I), Burroughs Wellcome Fund (C.B.W.), and Emergent Ventures Fast Grant (A.I., C.B.W). T.M. is supported by NIAID T32AI007019. B.I. is supported by NIAID 2T32AI007517-16 and K08AI163493. C.L. is a Pew Latin American Fellow. C.B.F.V. is supported by NWO Rubicon 019.181EN.004. C.B.W. is supported by K08AI128043. A.I. and A.M.P. are investigators of the Howard Hughes Medical Institute.

## Consortia (Yale SARS-CoV-2 Genomic Surveillance Initiative)

Tara Alpert, Anderson F. Brito, Rebecca Earnest, Joseph R. Fauver, Chaney C. Kalinich, Ketty Munyenyembe, Isabel M. Ott, Mary E. Petrone, Jessica Rothman, Anne E. Watkins

## Author Contributions

T.M. and A.I. designed the experiments; T.M., B.I., C.L., C.B.F.V., M.I.B., B.M., and H.D. performed experiments; T.M., B.I., C.L., C.B.F.V., N.D.G., and A.I. analyzed and interpreted results; O.F., B.M., M.L., C.W., M.L.L., N.D.G., and A.M.P. provided critical resources and reagents; T.M., B.I., and A.I. prepared the manuscript; A.I. secured funds and supervised the research. Authors from the Yale SARS-CoV-2 Genomic Surveillance Initiative contributed to sample screening, sample processing, SARS-CoV-2 genome sequencing, and data analysis.

## Corresponding Author

Correspondence and requests for materials should be addressed to Akiko Iwasaki.

---

**Extended Data Figure 1.**
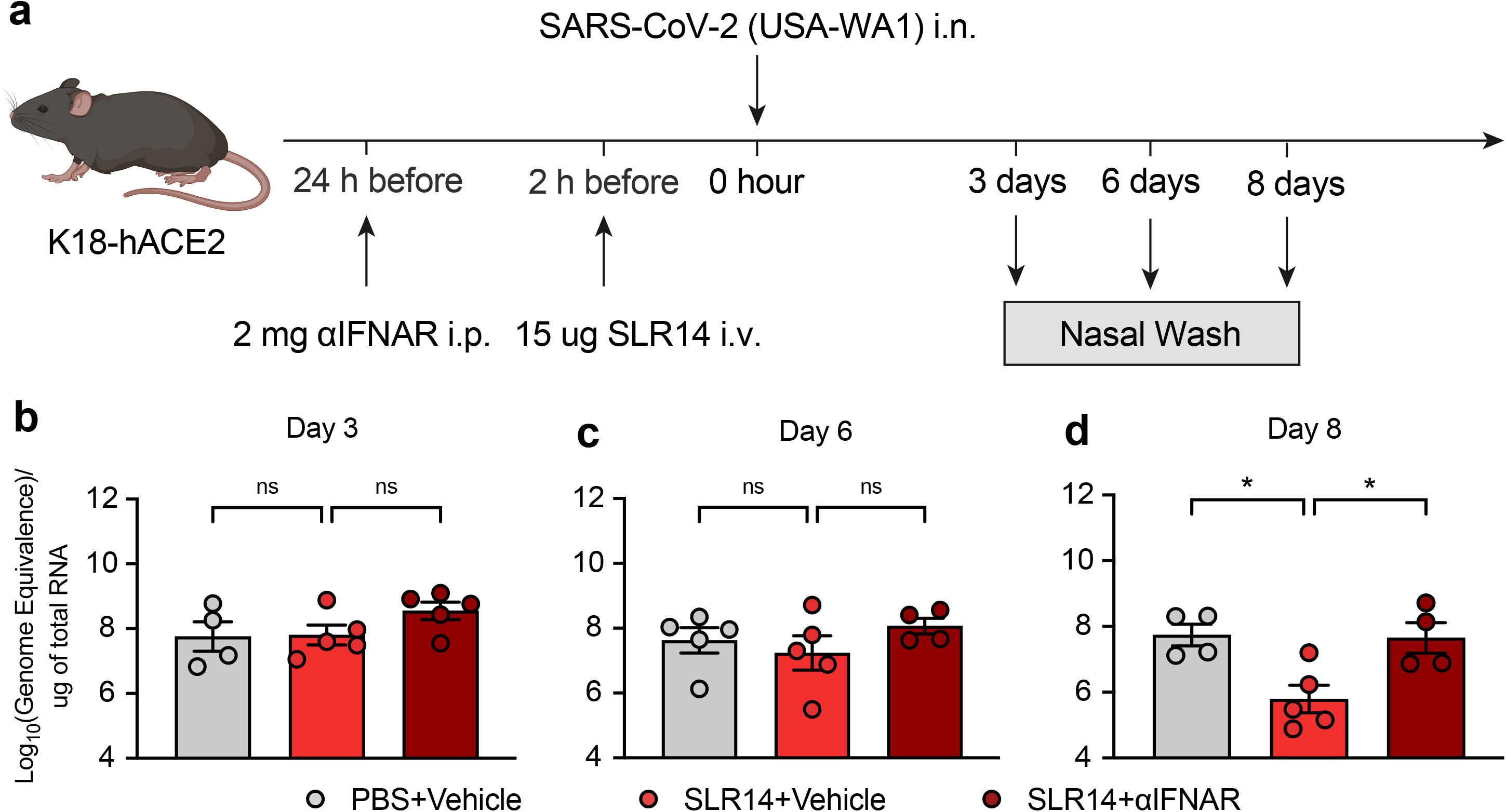
**a**, Experimental scheme: K18-hACE2 mice were intranasally infected with 10^3^ PFU SARS-CoV-2 (2019n-CoV/USA_WA1/2020). 2 hours before infection, 15 μg SLR14 or vehicle were intravenously administered. 24 hours before SLR14 injection, half of SLR14-treated mice was additionally given 2 mg anti-IFNAR antibodies. Nasal washes were collected for virological analysis 3, 6, and 8 DPI. **b-d**, Measurement of genomic viral RNA in the nasal wash 3, 6, and 8 DPI by RT-qPCR using the CDCN2 primer-probe set. Mean ± s.e.m.; Statistical significance was calculated by one-way ANOVA followed by Tukey correction (**b-d**); *P ≤ 0.05, **P ≤ 0.01, ***P ≤ 0.001, ****P ≤ 0.0001.

**Extended Data Figure 2.**
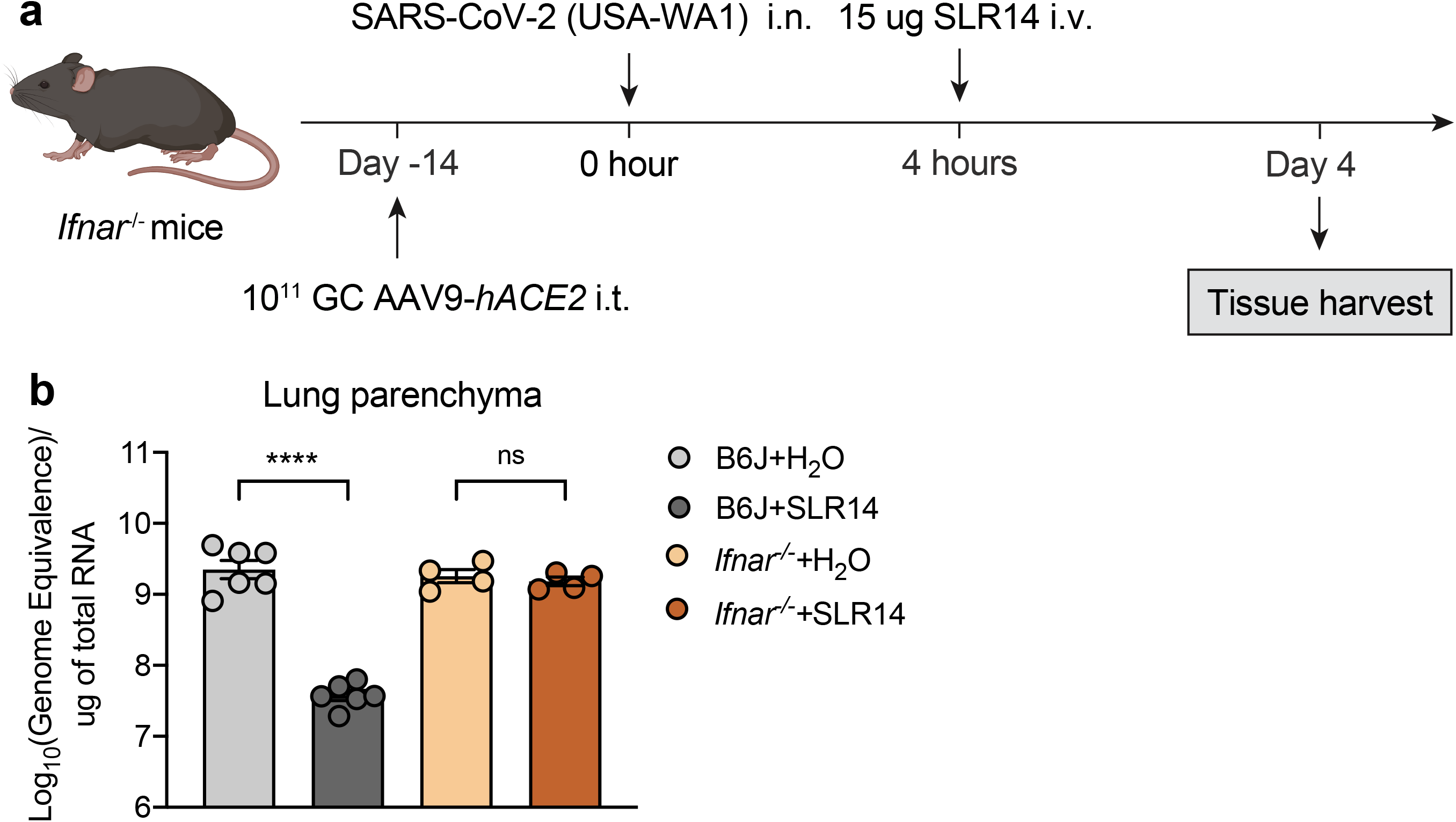
**a**, Experimental scheme: *Ifnar*^−/−^mice were intratracheally administered with 10^11^ genome copies of AAV9-*hACE2* and let rest for 2 weeks before intranasal infection with 10^6^ PFU SARS-CoV-2 (2019n-CoV/USA_WA1/2020). 15 μg SLR14 or vehicle were intravenously administered at 4 hours after infection. Lung tissues were collected for virological analysis 4 DPI. **b**, Measurement of genomic viral RNA 4 DPI by RT-qPCR using the CDCN2 primer-probe set. Mean ± s.e.m.; Statistical significance was calculated by one-way ANOVA followed by Tukey correction (**b**); *P ≤ 0.05, **P ≤ 0.01, ***P ≤ 0.001, ****P ≤ 0.0001.

**Extended Data Figure 3.**
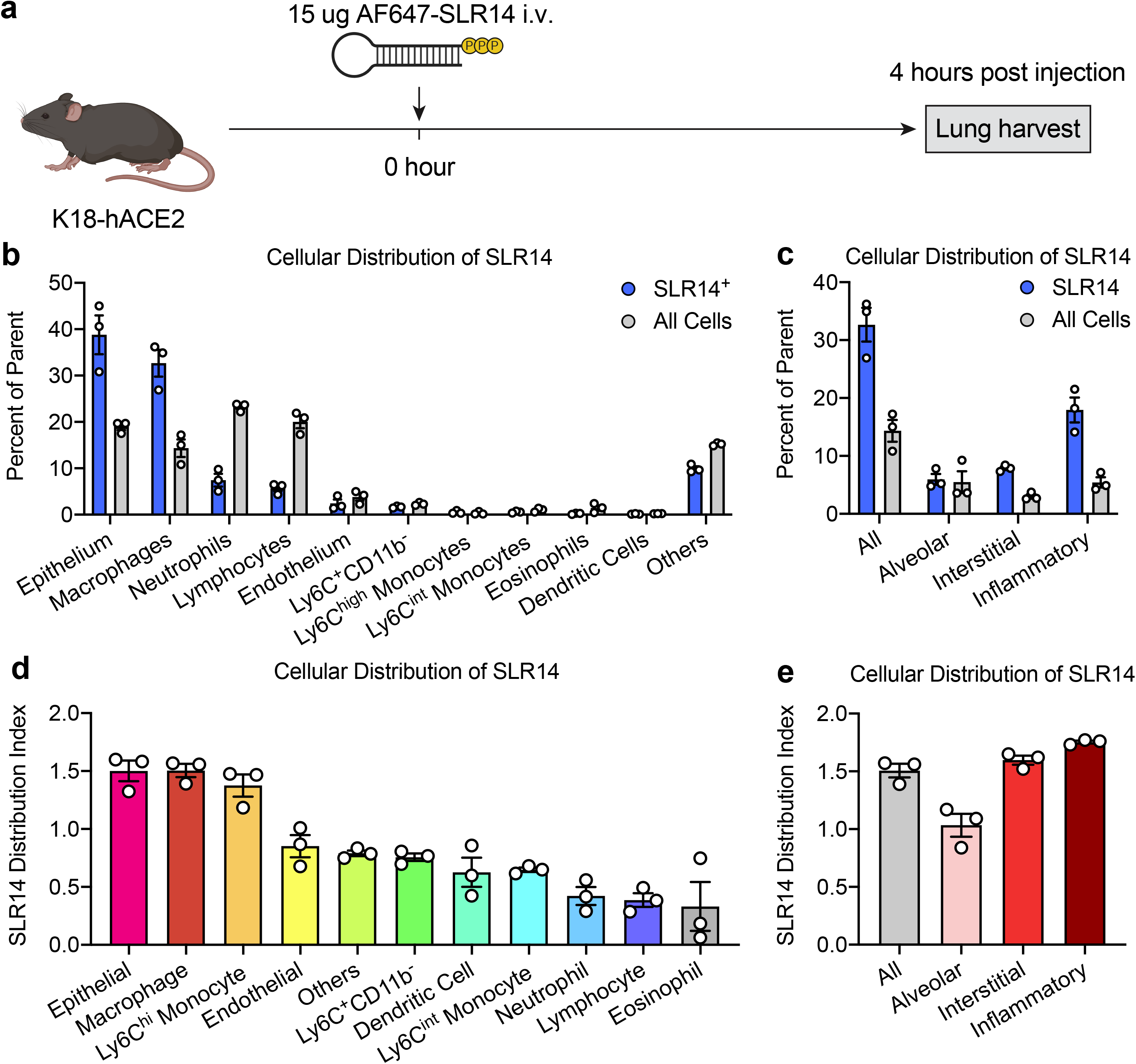
**a**, Experimental scheme: K18-hACE2 mice were i.v. injected with 15 μg AF647-conjugated SLR14 or vehicle. Lung tissues were collected for SLR14 uptake analysis by flow cytometry 4 hours post injection. Lung tissues from vehicle-injected controls were also collected as negative controls. **b**, Frequency of indicated immune and non-immune cell types among SLR14^+^ cells versus total lung cells. **c**, Frequency of indicated macrophage populations among SLR14^+^ cells or total lung cells. **d**, Distribution index (frequency of a given cell type in the SLR14^+^ compartment/frequency of all cells) of indicated immune and non-immune cell types. **e**, Distribution index of indicated macrophage populations.

**Extended Data Figure 4.**
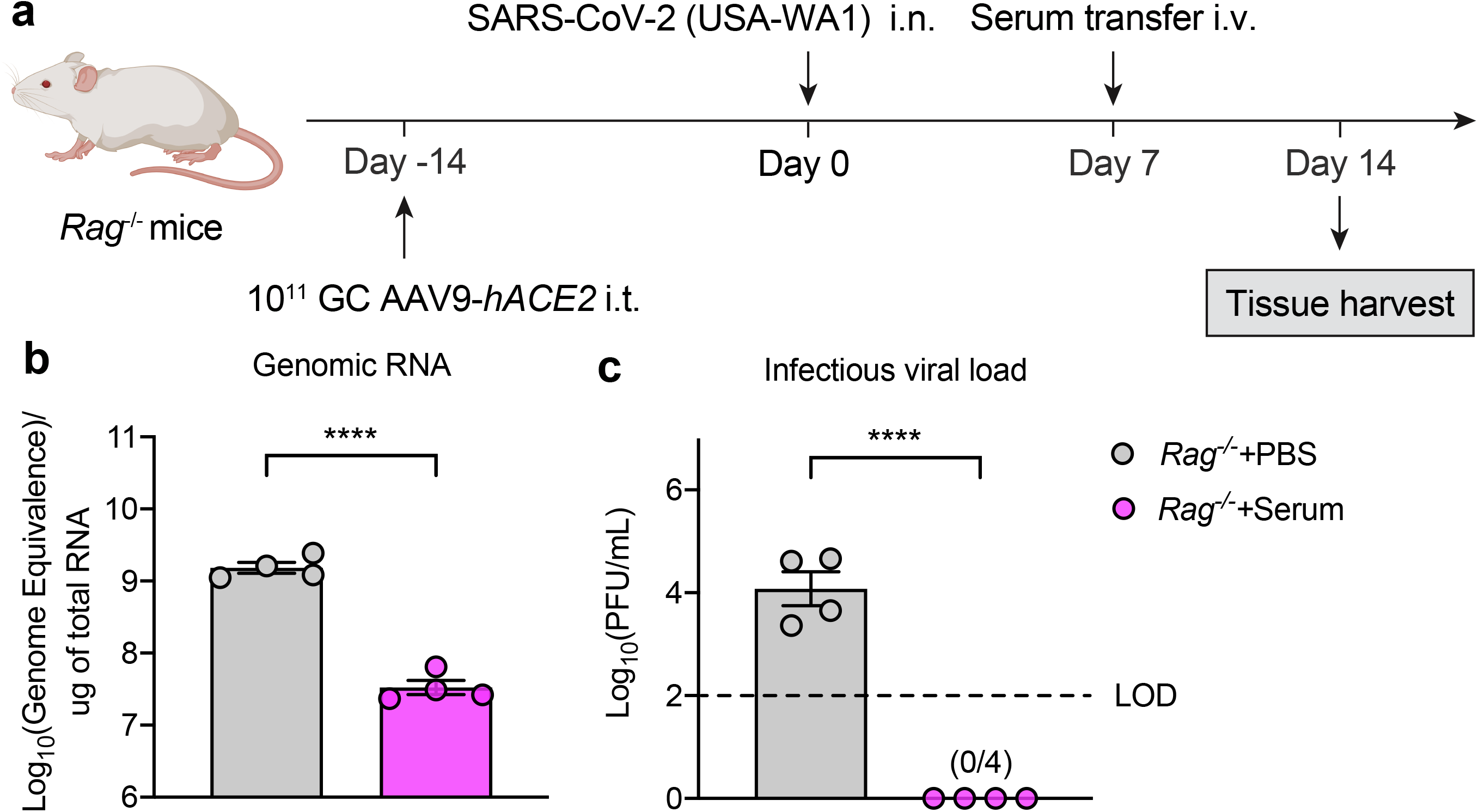
**a**, Experimental scheme: *Rag*^−/−^ mice were intratracheally administered with 10^11^ genome copies of AAV9-*hACE2* and let rest for 2 weeks before intranasal infection with 10^6^ PFU SARS-CoV-2 (2019n-CoV/USA_WA1/2020). 200 μL convalescent sera or PBS were intravenously administered 7 DPI. Lung tissues were collected for virological analysis 14 DPI. **b**, Measurement of genomic viral RNA in the lung 14 DPI by RT-qPCR. **c**, Measurement of infectious virus titer in the lung 14 DPI by plaque assay. Limit of detection (LOD) for plaque assay: 10^2^ PFU/mL. Mean ± s.e.m.; Statistical significance was calculated by one-way ANOVA followed by Tukey correction (**b,c**); *P ≤ 0.05, **P ≤ 0.01, ***P ≤ 0.001, ****P ≤ 0.0001. Data are pooled from two independent experiments.

**Extended Data Figure 5.**
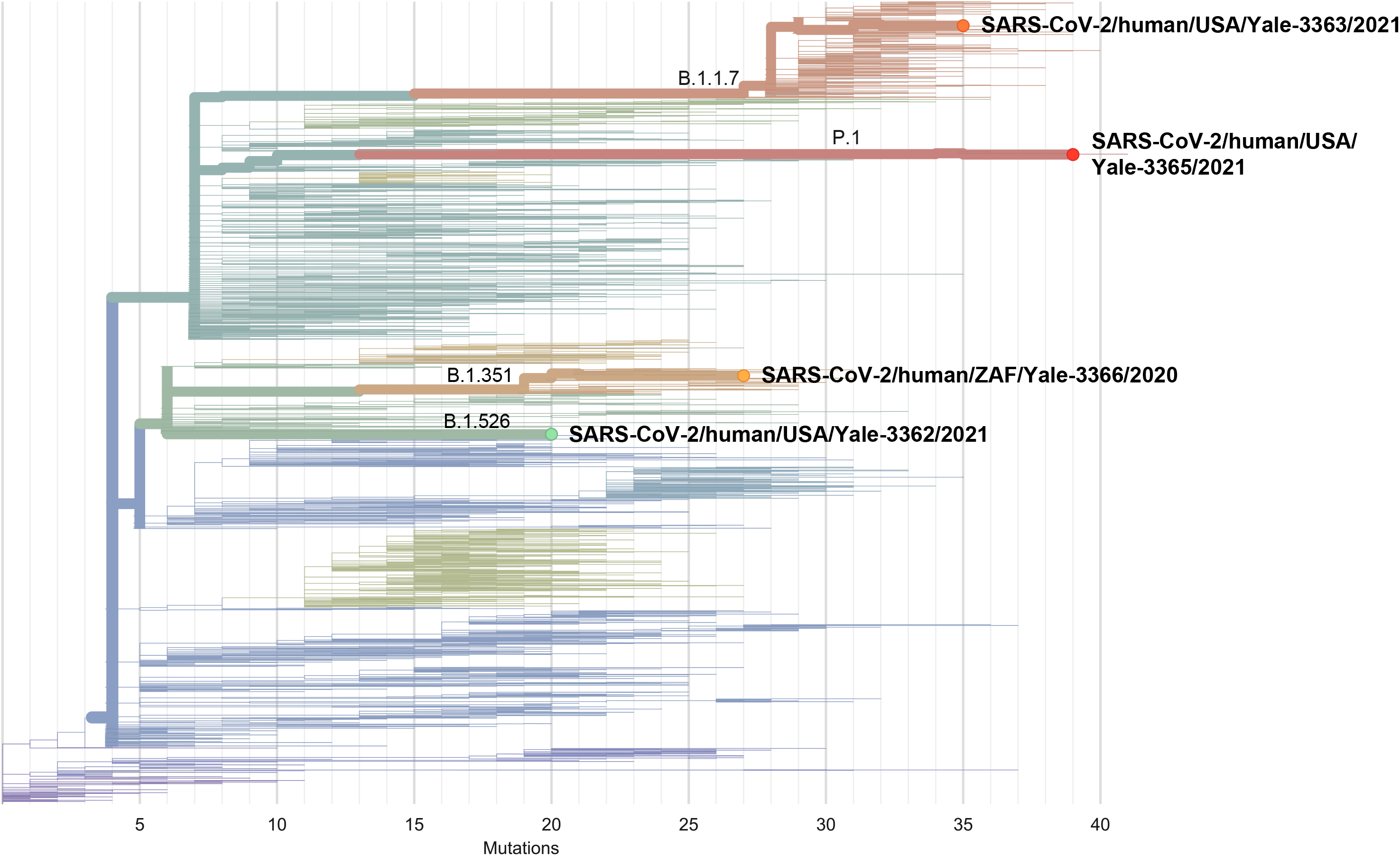
We used NextClade (https://clades.nextstrain.org/) to generate a phylogenetic tree to show the evolutionary relations between the four cultured viruses and other SARS-CoV-2 lineages. Branches are colored by Nextstrain clade, with branch labels indicating the Pango lineages, and highlighted are the four cultured viruses used in this study.

**Extended Data Figure 6.**
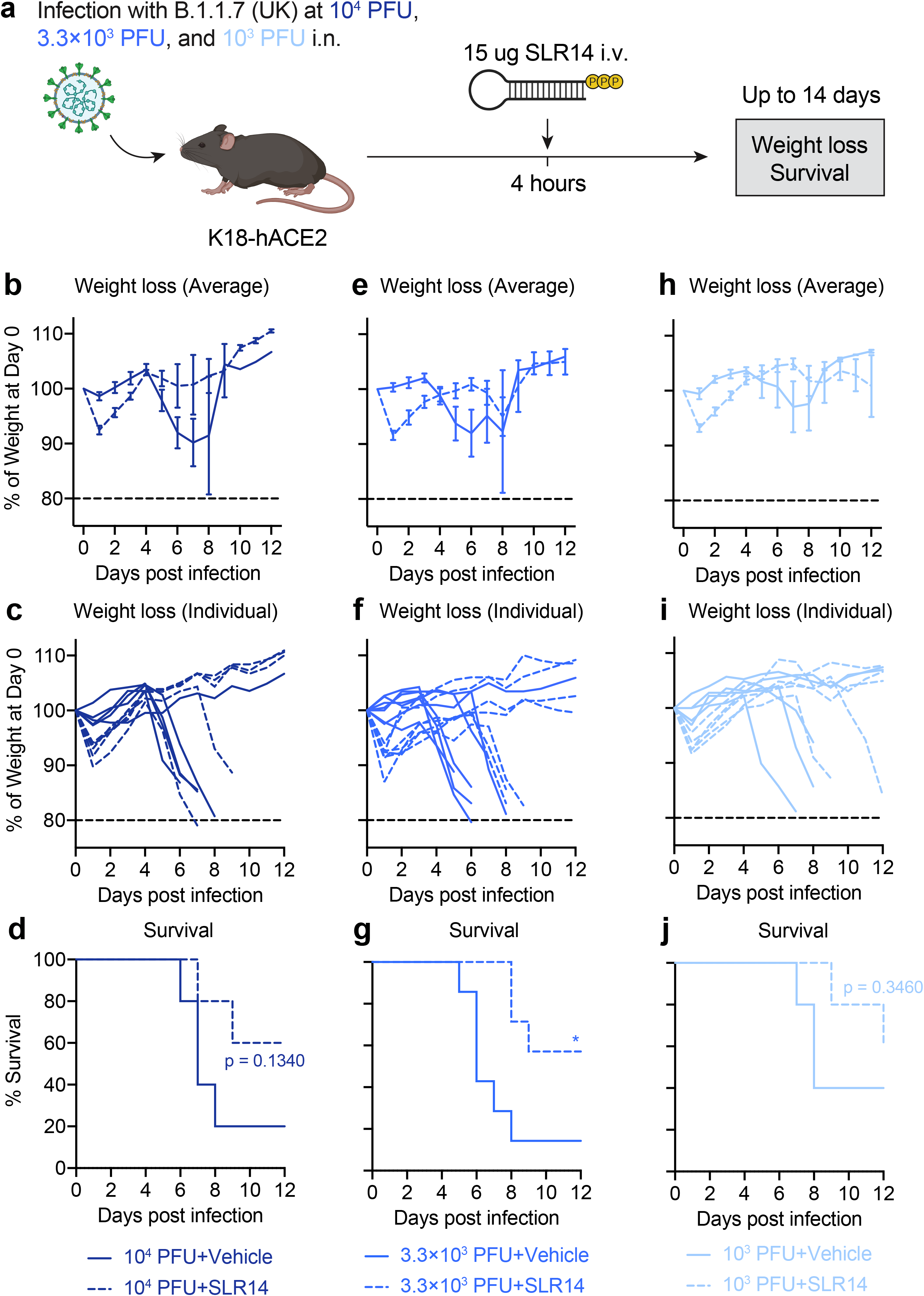
**a**, Experimental scheme: K18-hACE2 mice were intranasally infected with 10^4^, 3.3 × 10^3^, or 10^3^ PFU B.1.1.7 variant. 15 μg SLR14 or vehicle were intravenously administered at 4 hours after infection. Weight loss and survival were monitored daily up to 14 DPI. Death was recorded when mice were found dead in the cage, moribund, or at 80% of original body weight. **b-d**, Weight loss (**b,c**) and survival (**d**) of SLR14- and vehicle-treated K18-hACE2 mice from 1 to 14 DPI following infection with 10^4^ PFU B.1.1.7. **e-g**, Weight loss (**e,f**) and survival (**g**) of SLR14- and vehicle-treated K18-hACE2 mice from 1 to 14 DPI following infection with 3.3 × 10^3^ PFU B.1.1.7. **h-j**, Weight loss (**h,i**) and survival (**j**) of SLR14- and vehicle-treated K18-hACE2 mice from 1 to 14 DPI following infection with 10^3^ PFU B.1.1.7. Mean ± s.e.m.; Statistical significance was calculated by log-rank Mantel–Cox test (**d,g,j**); *P ≤ 0.05, **P ≤ 0.01, ***P ≤ 0.001, ****P ≤ 0.0001.

**Extended Data Figure 7.**
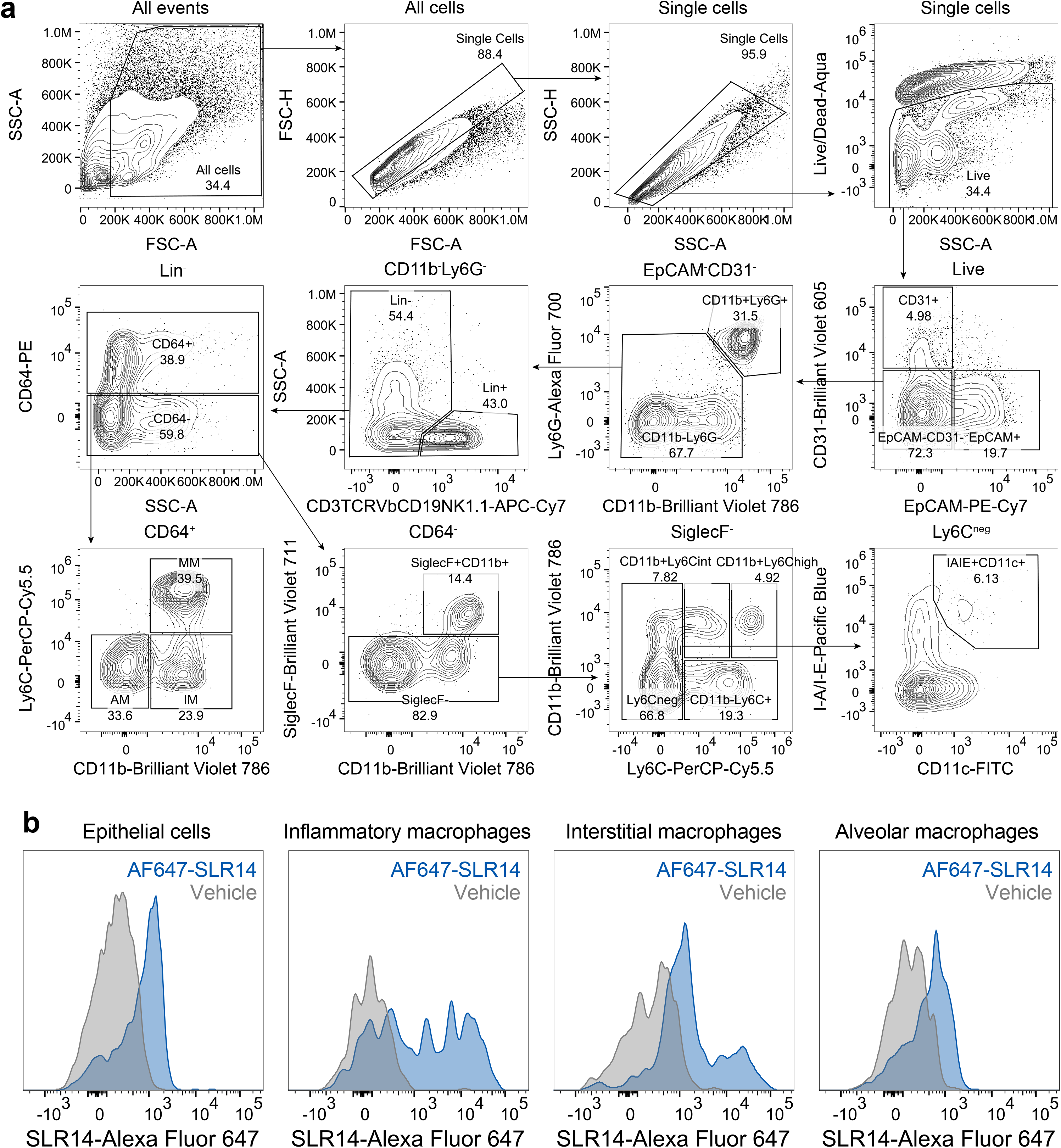
**a**, Gating strategies for identification of various immune and non-immune cell populations in the lung were used to generate Fig. 1j.k and **Extended Data Fig. 3b-e**. **b**, Histogram examples of SLR14 uptake by epithelial cells and different macrophage subsets. Lung tissues from vehicle-treated mice were included as negative controls.

