## Extended Data Table 1 for "A stem-loop RNA RIG-I agonist confers prophylactic and therapeutic protection against acute and chronic SARS-CoV-2 infection in mice"

**Extended Data Table 1: Amino acid changes identified in SARS-CoV-2 re-sequenced after virus isolation as compared to the reference genome (Accession MN908947)**.

Listed are amino acid substitutions and deletions for each of the genes. Letters indicate amino acids, numbers indicate amino acid positions, asterisks indicate stop codon mutations, and dashes indicate deletions. Underlined mutations are lineage defining.

| GenBank accession | **B.1.1.7**  MZ202178 | **B.1.351**  MZ202314 | **P.1**  MZ202306 | **B.1.526**  MZ201303 |
| --- | --- | --- | --- | --- |
| E |  | P71L |  |  |
| N | M1X  D3L  R203K  G204R  S235F | T205I | P80R  R203K  G204R | M1X  P199L  M234I |
| ORF1a | T1001I  P1213L  A1708D  I2230T  M2259I  S3675-  G3676-  F3677- | T265I  K1655N  K3353R  S3675-  G3676-  F3677- | S1188L  K1795Q  G2941S  S3675-  G3676-  F3677- | T265I  T2977I  L3201P  S3675-  G3676-  F3677- |
| ORF1b | P218L  P314L  A1432V | P314L | P314L  A1219S  E1264D | P314L  Q1011H  R1078C |
| ORF3a |  | Q57H  W131L  S171L | Q57H  S253P | P42L  Q57H |
| ORF7a |  | V93F |  |  |
| ORF8 | Q27*  R52I  K68*  Y73C | R115L | E92K | T11I |
| ORF9b |  |  | Q77E |  |
| S | H69-  V70-  Y144-  N501Y  A570D  D614G  P681H  T716I  S982A  D1118H | L18F  D80A  D215G  L242H  K417N  E484K  N501Y  D614G  Q677H  R682W  A701V  A243-  L244-  H245- | L18F  T20N  P26S  D138Y  R190S  K417T  E484K  N501Y  D614G  H655Y  T1027I  V1176F | L5F  T95I  D253G  E484K  D614G  A701V |

Abbreviations: E = envelope protein, N = nucleocapsid protein, ORF = open reading frame, S = spike protein.
